# Programmable DNA Assemblies Reconstitute Supramolecular Protein Function

**DOI:** 10.64898/2026.09.15.751908

**Authors:** Dhanush Gandavadi, Abhisek Dwivedy, Revathi Manoharaan, Seongmin Im, Saurabh Umrao, Chau Nguyen Minh Hoang, Kadmos Hammoud, Lauren Nguyen, Yang Zhao, Rohit Bhargava, Xing Wang

**Author notes:** Corresponding author: Xing Wang.

## Abstract

Biological processes are often regulated through reactions involving supramolecular protein complexes organized with nanoscale precision, such as the apoptosome that activates procaspase-9 through defined stoichiometry to induce apoptosis. Here we report designer CASCAD (<u>C</u>ARD <u>A</u>ssembly on <u>SCA</u>ffold <u>D</u>NA), a self-assembled DNA nanoplatform that spatially organizes caspase recruitment domains (CARDs), exhibiting catalytic intracellular caspase-9 activation. By functioning like a natural apoptosome, CASCAD enhances caspase-9 oligomerization and promotes valency-dependent enzymatic activation. Further, after incorporating targeting aptamers and membrane-interacting peptides for cellular uptake and cytosolic delivery, CASCAD has successfully reconstituted apoptosome function and downstream apoptotic signaling in *in-vitro* 2D and 3D cultures. Our molecular design paradigm establishes CASCAD as versatile and programmable DNA nanomaterial for reconstructing signaling pathways driven by diverse supramolecular complexes, including inflammasomes and myddosomes, providing a general chemical framework for engineering cellular signaling and directing cellular function and fate.

## INTRODUCTION

Supramolecular assemblies spatially organize signaling proteins with nanometer-scale precision to transmit, amplify and integrate information in living cells^1,2^. Across many intracellular pathways, signaling output is governed not only by the biochemical identity of signaling molecules, but also the spatial arrangement, stoichiometry, valency and proximity of interacting components within higher-order molecular complexes^3^. Such supramolecular organizing centers (SMOCs) function as control hubs that coordinate and catalyze signal initiation, amplification and propagation by locally concentrating signaling factors and regulating their collective interactions^4^. SMOCs are therefore fundamental determinants of cellular decision-making processes, including innate immune activation, inflammatory signaling and programmed cell death or apoptosis^5,6^.

The apoptosome is a heptameric SMOC that serves as the central signaling hub of the intrinsic apoptotic pathway, essential for maintaining cellular homeostasis^7^. In response to intracellular stressors such as DNA damage and oxidative stress, mitochondrial outer membrane permeabilization leads to the cytosolic release of cytochrome c^8^. Cytochrome c subsequently binds apoptotic protease-activating factor-1 (Apaf-1), triggering the assembly of the apoptosome that orchestrates caspase-9 catalysis ^9–11^. Within the apoptosome, procaspase-9 is recruited through homotypic caspase recruitment domain (CARD) interactions, resulting in nanoscale clustering and spatial confinement that promote oligomerization, initiating activation of caspase-9^12,13^. Activated caspase-9 subsequently drives the downstream caspase cascade, leading to apoptosis^14,15^. Importantly, the apoptosome demonstrates how intracellular signaling can emerge from the spatial organization of proteins into defined supramolecular architectures, where geometry, stoichiometry and proximity directly regulate enzyme catalysis^16,17^.

The ability to construct a synthetic SMOC capable of such intracellular signaling would provide powerful tools for both fundamental studies of spatial biochemistry and the development of programmable systems capable of controlling cell fate^18^. However, engineering artificial SMOCs that function intracellularly remains highly challenging^19^. Synthetic systems must not only recreate the nanoscale organization required for proximity-dependent catalysis, but also maintain structural programmability, biochemical stability, cellular uptake and cytosolic accessibility within the intracellular environment^20^. Although protein engineering and biomaterial-based approaches have enabled partial reconstruction of signaling assemblies, achieving precise and programmable control over intracellular SMOCs remains an immense and unresolved challenge^21–23^.

DNA nanotechnology offer a unique and versatile framework for engineering molecular organization with nanometer precision^24,25^. In particular, designer DNA nanostructures enable the programmable assembly of nanostructures with defined shape, size, and geometry, allowing functional moieties to be positioned with near-molecular accuracy and stoichiometry^26^. These properties have enabled DNA nanostructures to serve as scaffolds for organizing enzymes^27^, controlling multivalent interactions^28^, probing receptor signaling^29^, and studying geometry-dependent biological phenomena^30^. More broadly, DNA nanostructures have increasingly emerged as synthetic platforms for manipulating cellular processes through functional and spatial engineering. Despite these advances, the use of DNA nanotechnology to construct synthetic SMOCs that activate endogenous signaling pathways remains largely unexplored. Previous studies have primarily focused on extracellular receptor organization and *in vitro* enzymatic reconstitution^31–39^, which do not fully recapitulate the complexity of intracellular SMOCs. In the context of apoptotic signaling, synthetic reconstruction of apoptosome-like activity inside living cells remains particularly challenging because it requires the integration of programmable nanoscale protein organization with intracellular delivery, cytosolic catalytic functionality and biologically productive outcomes^40^.

Here we report designer CASCAD (<u>C</u>ARD <u>A</u>ssembly on <u>SCA</u>ffold <u>D</u>NA), a self-assembled DNA nanoplatform designed to carry CARDs for programing proximity-induced catalytic activation of caspase-9 in cells. By spatially organizing seven caspase recruitment domains within a defined nanoscale architecture to reconstitute key structural and functional features of natural apoptosome, we designed and synthesized CASCAD-7 that has efficiently promoted initiator caspase activation through CARD clustering. To maintain intracellular functionality, CASCAD-7 further incorporates targeting aptamers and membrane-interacting peptides that facilitate cellular uptake and cytosolic access respectively. We demonstrate that CASCAD-7 has successfully induced robust caspase-9 activation and apoptosis in cancer cells, reaching levels comparable to staurosporine, a prototypical bacteria-derived agent. CASCAD-7 activates apoptotic signaling through a programmable scaffold-mediated mechanism based on CARD clustering and proximity-induced caspase-9 activation. The results establish programmable DNA nanostructures as synthetic SMOCs capable of modulating endogenous cell-fate pathways.

## RESULTS AND DISCUSSION

### Design and characterization of CASCAD-7

DNA origami enables the precise spatial organization of biomolecules with nanometer-scale control, providing a programmable framework for reconstructing higher-order signaling assemblies. Inspired by the native human apoptosome structure (PDB: 5JUY)^41^, which exhibits a characteristic 7:4 stoichiometry of Apaf-1 to caspase-9, we sought to engineer a synthetic platform capable of recapitulating the nanoscale organization underlying procaspase-9 activation (**Fig. 1a**). The apoptosome contains a central disk-like assembly of seven Apaf-1 CARDs that plays a key role in recruiting procaspase-9 through homotypic CARD (Apaf-1) - CARD (Caspase-9) interactions (**Fig. 1b**)^42–44^. Structural analyses of apoptosome-associated CARD organization together with previous studies on proximity-induced caspase-9 activation demonstrated that intermolecular separations in the ∼4-6 nm range are favorable for caspase interactions and maximal enzymatic activation^45^. Based on these observations, we designed CASCAD, with neighboring CARDs separated by ∼6 nm to promote efficient caspase clustering while preserving sufficient conformational flexibility for productive CARD-CARD interactions and minimizing steric crowding within the scaffold.

**Fig. 1:**
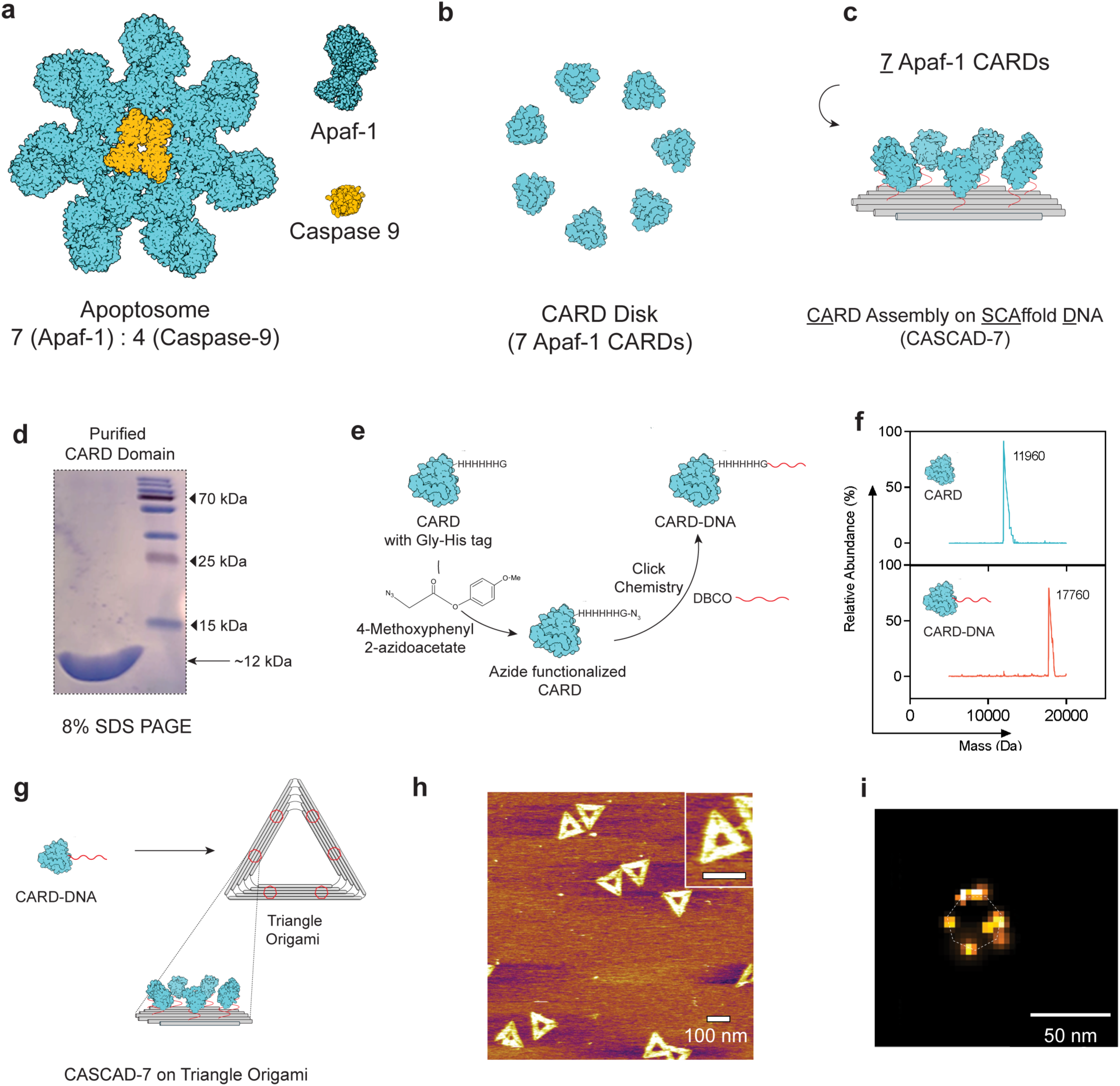
Design and characterization of CASCAD-7. **a.** Structural model of the native apoptosome illustrating the 7:4 stoichiometry between Apaf-1 and caspase-9. **b.** Schematic representation of the heptameric Apaf-1 CARD disk organization responsible for procaspase-9 recruitment and activation. **c.** Design concept of CASCAD-7 (<u>C</u>ARD <u>A</u>ssembly on <u>SCA</u>ffold <u>D</u>NA), in which seven CARD domains are spatially organized on a DNA origami scaffold. **d,** SDS-PAGE analysis of purified recombinant CARD domain showing a dominant band at ∼12 kDa. **e.** Site-specific functionalization and conjugation strategy for generating CARD-DNA conjugates using 4-Methoxyphenyl 2-azidoacetate and copper-free click conjugation with DBCO-modified oligonucleotides. **f.** Mass spectrometry analysis confirming successful formation of CARD-DNA conjugates through the expected molecular weight shift relative to unconjugated CARD protein. **g.** Assembly of CARD-DNA conjugates onto triangular DNA origami scaffolds to generate CASCAD-7 nanostructures with spatially organized CARD presentation. **h.** AFM image of assembled triangular DNA origami structures following CASCAD-7 assembly. Scale bar, 100 nm. **i.** Super-resolution fluorescence image showing localized heptameric clustering on CASCAD-7. Scale bar, 50 nm.

To reconstruct this architecture synthetically, we designed CASCAD-7, in which seven CARD domains are spatially organized on a DNA origami scaffold to mimic the native heptameric arrangement of the apoptosome CARD disk (**Fig. 1c**). The CARD domain was recombinantly expressed and purified to homogeneity, yielding a dominant species at ∼12 kDa as confirmed by SDS-PAGE analysis (**Fig. 1d** and **Supplementary Fig. 1**). To enable site-specific assembly onto DNA nanostructures, purified CARD proteins were functionalized with azide groups using 4-Methoxyphenyl 2-azidoacetate and subsequently conjugated to DBCO-modified oligonucleotides using copper-free click chemistry (**Fig. 1e**)^46^. Optimization of the conjugation conditions minimized multi-acylated species and improved formation of discrete CARD-DNA conjugates (**Supplementary Fig. 2**). Successful conjugation was confirmed by mass spectrometry through the expected molecular-weight shift of the CARD-DNA product (**Fig. 1f**).

We selected a triangular-shaped DNA origami (or called “Triangle Origami” herein) owing to its open, unbendable two-dimensional (2D) geometry, which provides much higher 2D structural rigidity than other 2D scaffolds, while minimizing steric crowding due to the central hollow region^47,48^. Previous studies have also shown that Triangle Origami structures exhibit efficient cellular internalization and favorable intracellular delivery properties^49–51^. To maximize multivalent CARD presentation and spatial organization, the Triangle Origami scaffold was engineered with six distinct regions for assembling CASCAD-7 corresponding to a total of 42 addressable hybridization sites, enabling controlled localization of CARD-functionalized oligonucleotides across the nanostructure (**Fig. 1g** and **Supplementary Fig. 3a**). Fluorescence gel analysis demonstrated that a triangle origami-to-Cy5 oligonucleotide ratio of approximately 1:42 resulted in near-saturating occupancy, consistent with the designed 42-site addressable sites on the scaffold (**Supplementary Fig. 3b**).

CARD-DNA conjugates were subsequently assembled onto DNA origami to generate the complete CASCAD-7 nanostructure and gel shift assay confirmed the binding of CARD-DNA to Triangle Origami (**Supplementary Fig. 4**). Atomic force microscopy (AFM) imaging demonstrated formation of intact Triangular Origami particles with localized regions of CARD domains (**Fig. 1h** and **Supplementary Fig. 5**). As individual CARD domains were below the direct lateral resolution limit of AFM, super-resolution fluorescence imaging was further used to validate nanoscale localization of individual CASCAD-7 cluster on the origami scaffold (**Fig. 1i**). Quantitative occupancy analysis revealed an average loading of 6.7 ± 1.0 CARD assemblies per cluster on origami (**Supplementary Fig. 6**), consistent with efficient heptameric organization. Together, these results establish CASCAD-7 that reconstructs key nanoscale features of the CARD disk from the native apoptosome.

### CASCAD-7 promotes multivalent recruitment and proximity-driven activation of caspase-9

A defining feature of the native apoptosome is its ability to locally concentrate procaspase-9 through multivalent CARD-CARD interactions, thereby increasing the effective local concentration of caspase-9 and promoting proximity-induced activation^17,52^. We therefore sought to determine whether the CASCAD-7 enhances caspase-9 recruitment and catalytic activation.

To first evaluate binding, we quantified the association of caspase-9 with monomeric CARD-DNA conjugate and fully assembled CASCAD-7 nanostructures using surface plasmon resonance (SPR) (**Fig. 2a**). Purified caspase-9 was confirmed by SDS-PAGE prior to binding measurements, revealing the expected p35 and p10 subunits following processing (**Supplementary Fig. 7a**). Monomeric CARD-DNA displayed relatively weak binding toward immobilized caspase-9, with a dissociation constant (K_D_) of 361.9 nM (**Fig. 2b** and **Supplementary Fig. 7b**). In contrast, CASCAD-7 exhibited substantially enhanced binding affinity, yielding a K_D_ of 1.89 nM (**Fig. 2c** and **Supplementary Fig. 7c**), corresponding to an approximately 190-fold increase in apparent affinity relative to monomeric CARD presentation. Kinetic fitting further revealed a pronounced increase in the association rate constant (K_on_) for CASCAD-7 while maintaining a comparable dissociation rate (K_off_), consistent with multivalent avidity arising from clustered CARD presentation on the DNA scaffold (**Supplementary Fig. 7d**). These results demonstrate that nanoscale organization of CARD domains within CASCAD-7 strongly enhances caspase-9 recruitment through cooperative multivalent interactions.

**Fig. 2:**
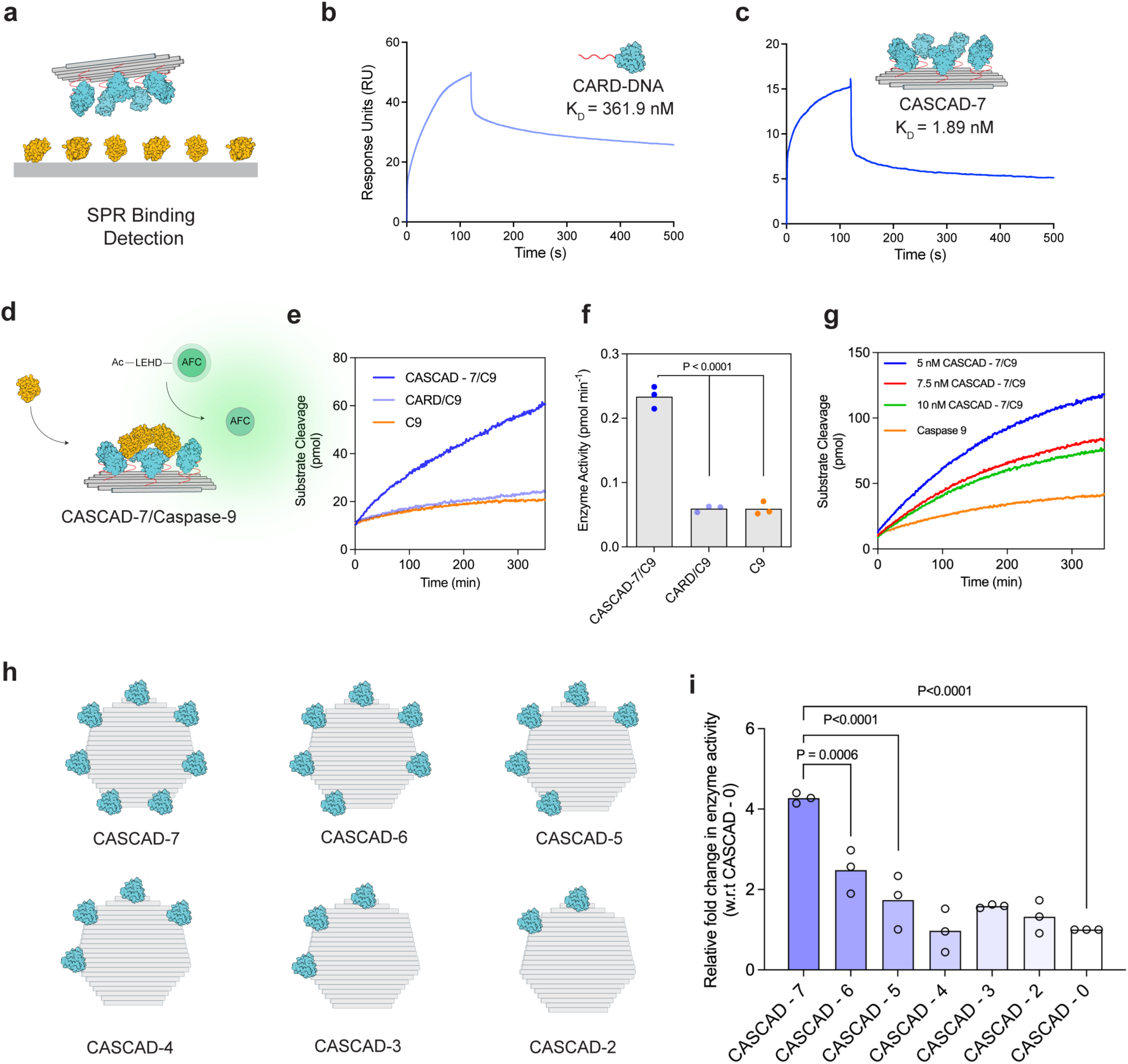
CASCAD-7 enables enhanced caspase-9 binding and activation. **a.** Schematic of surface plasmon resonance (SPR) assay used to measure binding between Caspase-9 (C9) and CASCAD-7. Representative SPR sensorgrams showing binding of monomeric CARD-DNA conjugates (**b**) and CASCAD-7 assemblies (**c**) to immobilized caspase-9. CASCAD-7 exhibited substantially enhanced binding affinity relative to monomeric CARD-DNA. **d.** Schematic illustrating recruitment and activation of C9 on the CASCAD-7 scaffold, resulting in cleavage of the fluorogenic LEHD-AFC substrate. **e.** Time-dependent substrate cleavage assays comparing C9 activity in the presence of CASCAD-7/C9 (1 nM CASCAD-7, 25 nM C9), monomeric CARD-DNA/C9 (42 nM CARD-DNA, 25 nM C9) and caspase-9 alone (25 nM). **f.** Quantification of enzymatic activity demonstrating enhanced caspase-9 activation mediated by CASCAD-7 relative to controls. **g.** Caspase-9 substrate cleavage kinetics measured at increasing CASCAD-7 concentrations (5 to 10 nM) while maintaining constant C9 concentration (125 nM). **h.** Schematic representation of CASCAD variants with decreasing CARD valency (CASCAD-7 to CASCAD-2) used to investigate the effect of multivalency on caspase-9 activation. **i.** Relative caspase-9 enzymatic activity as a function of CARD valency. Higher-order CARD assemblies produced progressively enhanced enzymatic activation compared to lower-valency and scaffold-only controls. Data are presented as mean ± s.e.m for *n* = 3 biologically independent samples. P values determined by one-way ANOVA followed by Dunnett’s multiple comparisons test.

We next investigated whether enhanced recruitment translated into functional caspase catalytic activation. To this end, caspase-9 was incubated with CASCAD-7 and enzymatic activity was quantified using the fluorogenic substrate LEHD-AFC, in which substrate cleavage results in release of fluorescent AFC (**Fig. 2d**). Fluorescence intensities were converted to picomoles of released AFC using a standard calibration curve to quantify enzymatic activity (**Supplementary Fig. 8a-b)**. CASCAD-7 induced robust caspase-9 activation compared to both monomeric CARD-DNA conjugates and free caspase-9 controls (**Fig. 2e**). Quantitative analysis of caspase-9 activation revealed that CASCAD-7 induced an ∼4 - 4.5-fold increase in enzymatic activity relative to basal caspase-9 levels, broadly aligning with previous observations from DNA-tethered caspase-9 systems^53,54^, whereas monomeric CARD–DNA conjugates produced only minimal enhancement (**Fig. 2f**). These findings indicate that spatial clustering of CARDs on the origami scaffold efficiently promoted caspase activation.

To further probe the relationship between CASCAD stoichiometry and signaling output, we varied the concentration of CASCAD-7 while maintaining a constant concentration of caspase-9 (**Fig. 2g**). Maximal enzymatic activity was observed at a CASCAD-7 concentration corresponding to an approximate 1:4 CASCAD-7 to caspase-9 ratio, consistent with the native apoptosome stoichiometry, whereas increasing scaffold concentrations progressively reduced catalytic efficiency. This behavior is consistent with a proximity-driven activation mechanism in which optimal signaling requires productive recruitment of multiple caspase-9 molecules onto individual scaffolds. At elevated CASCAD-7 concentrations, caspase-9 molecules become distributed across separate nanostructures, thereby reducing intermolecular dimerization and productive catalytic assembly, a phenomenon previously observed in scaffold-mediated signaling systems in which excess scaffold abundance reduces optimal signaling efficiency^55,56^. These observations further support the importance of optimal ratio for scaffold-mediated spatial confinement in regulating caspase catalysis.

Since the native apoptosome contains a heptameric CARD assembly, we next sought to directly examine the contribution of CARD valency to caspase-9 activation. We therefore generated a palette of CASCAD variants containing progressively reduced numbers of CARD domains, ranging from seven to zero organized CARD sites (CASCAD-7 to CASCAD-0) (**Fig. 2h**). Catalytic profiles revealed a clear valency-dependent activation profile, in which higher-order CARD assemblies produced progressively stronger caspase-9 activation (**Fig. 2i** and **Supplementary Fig. 8c**). Notably, the transition from lower-order assemblies to the heptameric configuration resulted in an enhanced response, suggesting that higher-order clustering promotes cooperative proximity-induced activation.

### CASCAD-7 platform enables targeted intracellular activation of caspase-9

To extend CASCAD-7 toward intracellular studies, we selected the human erythroleukemia cell line HEL92.1.7 as a disease-relevant model for evaluating programmable apoptotic activation. They exhibit dysregulated apoptotic signaling, including impaired initiator caspase catalysis and apoptosome dysfunction, which contribute to uncontrolled proliferation and therapeutic resistance^57,58^. We hypothesized that successful activation of endogenous caspase signaling in HEL92.1.7 cells would provide strong evidence that CASCAD-7 can function as synthetic intracellular signaling assembly capable of reactivating apoptosis.

To enable intracellular delivery and activation, we engineered a functionalized nanostructure capable of targeted cellular uptake and cytosolic delivery. CASCAD-7 was modified with leukemia targeting aptamers together with hemagglutinin (HA)-derived membrane-interacting peptides to facilitate endosomal escape following internalization (**Fig. 3a** and **Supplementary Fig. 9a**). The leukemia-targeting aptamer was previously validated by our group for selective recognition and uptake in leukemia cells^59,60^, while the HA peptide has been shown in earlier studies to enhance endosomal escape of triangular DNA origami nanostructures through reduced colocalization^50,61^. Successful conjugation of the HA peptide to DNA and assembly of the fully functionalized CASCAD-7 nanostructure incorporating both HA peptides and targeting aptamers were confirmed by agarose gel electrophoresis (**Supplementary Fig. 9 b-c**).

**Fig. 3:**
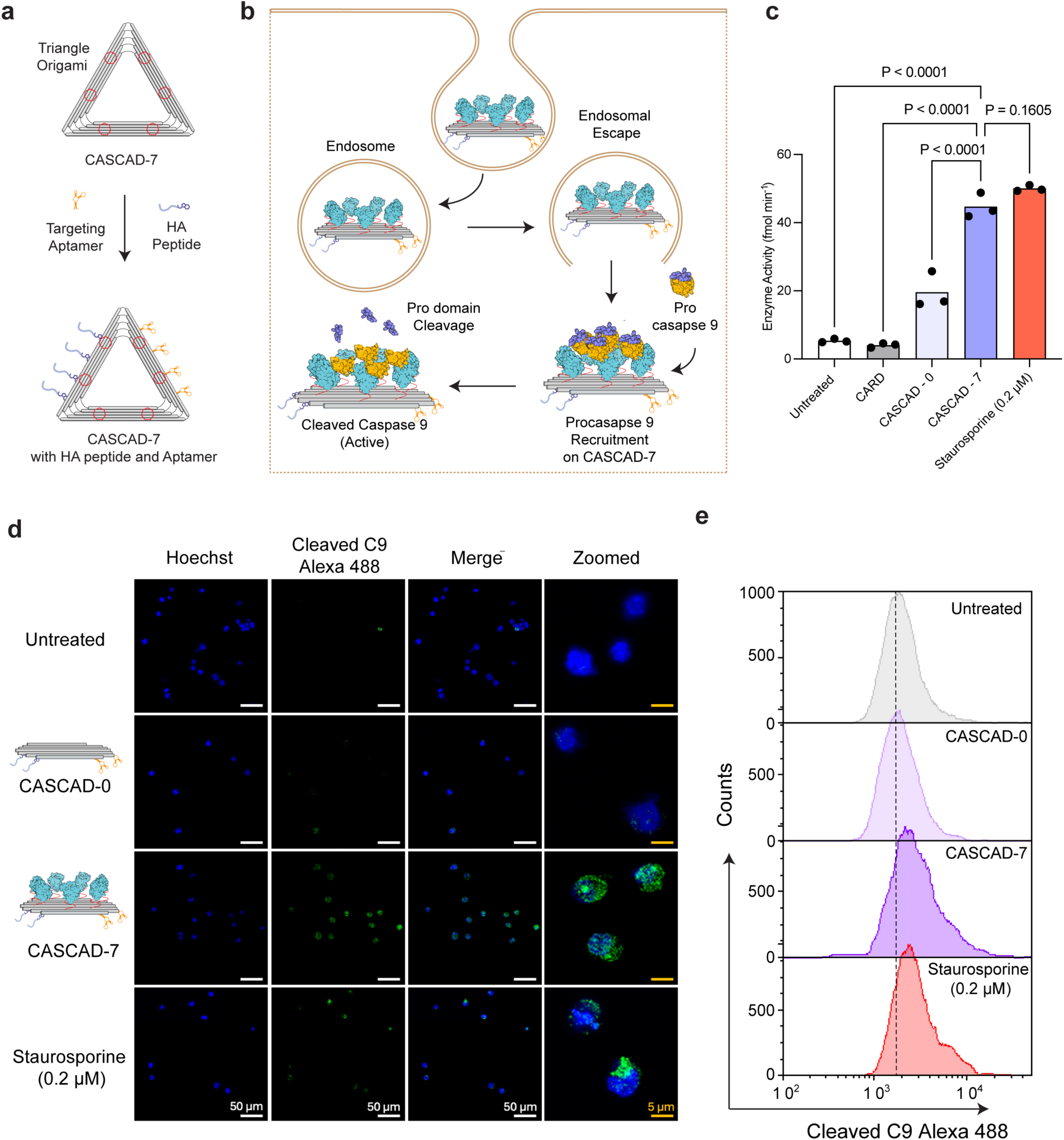
CASCAD-7 platform enables targeted intracellular activation of caspase-9. **a**. Schematic of triangular CASCAD-7 functionalized with HA peptide for membrane interaction and H9 aptamer for targeting. Agarose gel electrophoresis confirms successful assembly and functionalization of CASCAD-7. **b**. Proposed mechanism of action: CASCAD-7 undergoes selective cellular uptake, escapes the endosome using HA peptide, and recruits procaspase-9, facilitating proximity-induced activation through prodomain cleavage. **c**. Quantification of caspase-9 enzymatic activity across conditions shows significant activation in CASCAD-7-treated samples compared to controls. **d**. Confocal microscopy images showing cleaved caspase-9 (green) and nuclear staining (Hoechst, blue). CASCAD-7 treatment induces robust intracellular caspase-9 activation relative to controls. Scale bars: 50 μm (main), 5 μm (zoomed). **e**. Flow cytometry histograms of cleaved caspase-9 (Alexa 488) signal across treatment groups, demonstrating increased activation with CASCAD-7. Data are presented as mean ± s.e.m for *n* = 3 biologically independent samples. P values determined by one-way ANOVA followed by Dunnett’s multiple comparisons test.

Mechanistically, CASCAD-7 is designed to enter cells through receptor-mediated endocytosis followed by HA peptide-mediated destabilization of endosomal membranes, enabling release of the scaffold into the cytosol (**Fig. 3b**). Following cytosolic exposure, CASCAD-7 recruits endogenous procaspase-9 molecules onto its multivalent CARD platform through homotypic CARD-CARD interactions. Spatial confinement of procaspase-9 promotes proximity-induced dimerization and prodomain cleavage, generating catalytically active caspase-9 that subsequently initiates downstream apoptosis. As an initial measure of intracellular functionality, we quantified bulk caspase-9 enzymatic activity from cell lystates following treatment under different conditions (**Fig. 3c**). CASCAD-7 induced a substantial increase in caspase-9 activity relative to untreated cells, CARD-only controls and the non-functionalized scaffold (CASCAD-0). Notably, the magnitude of activation approached levels observed with the potent apoptosis inducer staurosporine (0.2 μM), highlighting CASCAD-7’s efficiency in driving intracellular caspase activation. In contrast, minimal activation was observed in control conditions, indicating that neither free CARD domains nor the origami scaffold alone are sufficient to induce efficient signaling without organized intracellular assembly.

To extend these findings beyond bulk lysate measurements and resolve caspase-9 activation at the single-cell level, we next examined intracellular caspase-9 cleavage as a readout of activation using confocal microscopy (**Fig. 3d**). Cells treated with CASCAD-7 for 12 hours displayed strong punctate and diffuse fluorescence corresponding to cleaved caspase-9, whereas untreated cells and control conditions exhibited minimal signal. Cleaved caspase-9 was observed throughout the cytosol and in perinuclear regions, consistent with successful cytosolic delivery of the nanostructure following endosomal escape. Higher-magnification images revealed localized clusters of cleaved caspase-9, potentially corresponding to regions where multiple procaspase-9 molecules are recruited onto individual CASCAD-7 scaffolds. Interestingly, caspase activation induced by CASCAD-7 appeared broadly throughout the cell, whereas staurosporine-treated cells exhibited more spatially localized activation. This difference may reflect the ability of intracellularly distributed CASCAD-7 to access and recruit procaspase-9 throughout the cytosol. These observations support a model in which nanoscale spatial organization enhances intracellular caspase activation through engineered proximity-induced assembly. Quantitative analysis of confocal fluorescence intensity further demonstrated substantially elevated caspase-9 cleavage signal in CASCAD-7-treated cells relative to untreated and control conditions (**Supplementary Fig. 10**). Interestingly, the fluorescence intensity observed for CASCAD-7 exceeded that of the staurosporine-treated positive control. Because staurosporine treatment resulted in extensive cell death and reduced cell density during imaging, these measurements may underestimate the total extent of apoptotic activation under these conditions. We therefore performed flow cytometry analysis to quantify cleaved caspase-9 activation across the entire cell population and more accurately compare apoptotic responses between treatment groups.

Flow cytometric analysis of cleaved caspase-9 further supported these observations (**Fig. 3e** and **Supplementary Fig. 11a**). Cells treated with CASCAD-7 for 12 hours exhibited a pronounced shift in fluorescence intensity relative to untreated cells and control conditions, indicating elevated intracellular levels of cleaved caspase-9 across the cell population. Quantification of peak fluorescence intensity further confirmed robust intracellular caspase-9 cleavage following CASCAD-7 treatment, showing substantially higher fluorescence intensity compared to CARD-only and CASCAD-0 controls and reaching approximately 80% of the signal observed in staurosporine-treated cells. (**Supplementary Fig. 11b**). We also observed a broader fluorescence distribution observed following staurosporine treatment likely reflects increased heterogeneity in apoptotic progression within the population, consistent with widespread and asynchronous activation of cell death pathways^62^.

### CASCAD-7 induces apoptotic signaling through activation of the caspase cascade

As we demonstrated CASCAD-7 promotes intracellular activation of caspase-9, we next investigated whether this caspase-9 activation propagates through the downstream apoptotic cascade to induce functional cell death. Mechanistically, recruitment of procaspase-9 onto the CASCAD-7 is expected to promote proximity-induced activation and proteolytic processing of caspase-9, which subsequently cleaves and activates caspase-3, ultimately driving apoptotic cell death (**Fig. 4a**).

**Fig. 4:**
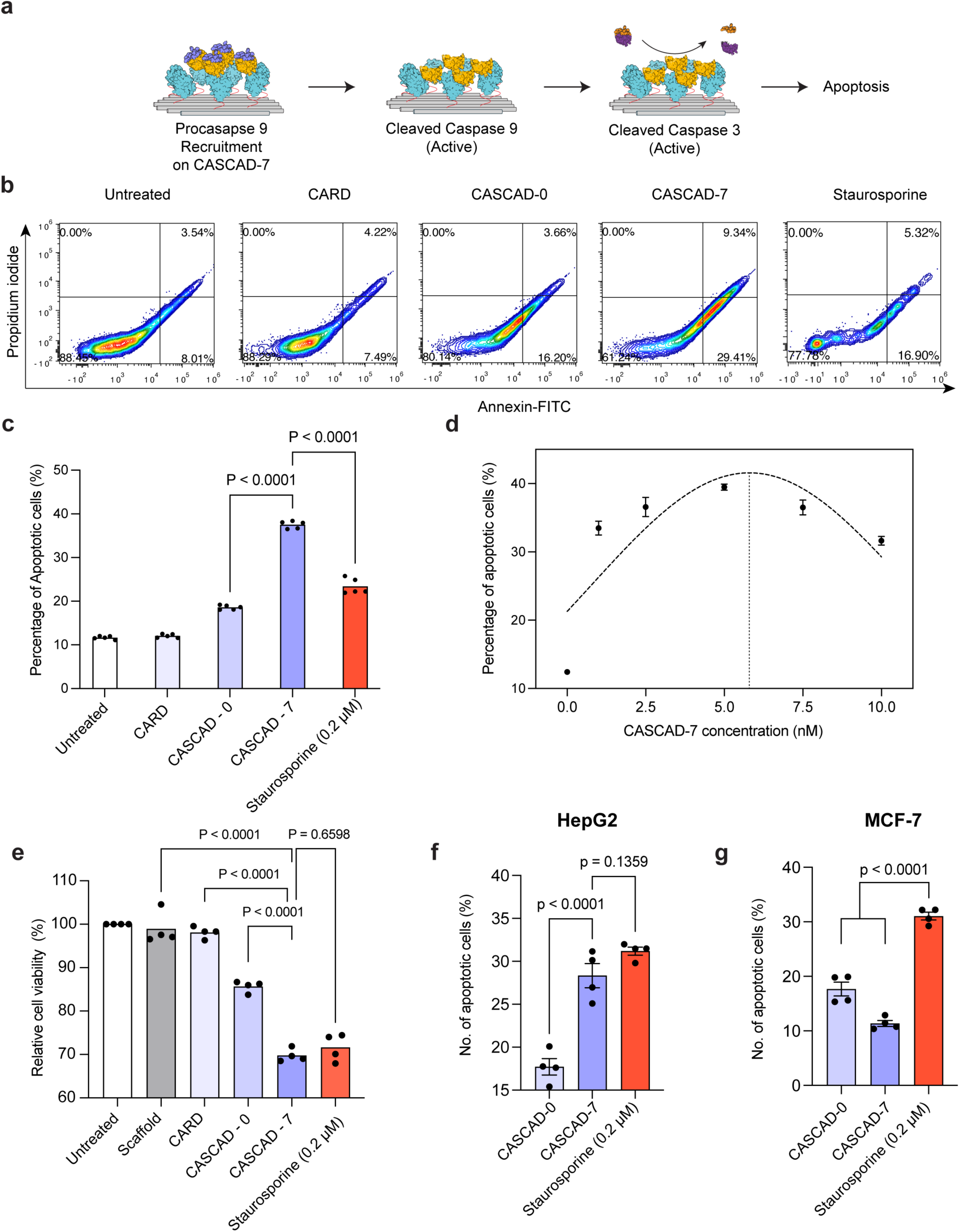
CASCAD-7 activates downstream apoptotic signaling and induces programmed cell death. **a**. Schematic illustrating proximity-induced activation of procaspase-9 on CASCAD-7, leading to downstream activation of executioner caspases and apoptosis. **b**. Representative Annexin V-FITC and propidium iodide (PI) flow cytometry contour plots of HEL92.1.7 cells following treatment with CARD, CASCAD-0, CASCAD-7 or staurosporine (0.2 µM). **c**. Quantification of apoptotic cell populations from Annexin V/PI analysis showing significantly increased apoptosis following CASCAD-7 treatment compared to untreated, CARD-only and CASCAD-0 controls. **d**. Concentration-dependent apoptotic response following treatment with increasing concentrations of CASCAD-7. **e**. Relative cell viability following treatment under indicated conditions. CASCAD-7 significantly reduced cell viability relative to untreated cells and control nanostructures, approaching levels observed with staurosporine treatment. **f & g.** Quantification of apoptotic cell populations in HepG2 (f) and MCF-7 (Caspase-3 Deficient) (g) cells. Data are presented as mean (**c,e**) or mean ± s.e.m (**d,f,g**) for *n* = 5 (**c,d**) *n* = 4 (**e,f,g**) biologically independent samples. P values determined by one-way ANOVA followed by Dunnett’s multiple comparisons test. Data are presented as mean or mean ± s.e.m for *n* = 4 or 5 biologically independent samples. P values determined by one-way ANOVA followed by Dunnett’s multiple comparisons test.

To quantify apoptosis, HEL92.1.7 leukemia cells were treated with CARD-only controls, CASCAD-0, CASCAD-7 or staurosporine as a positive apoptosis control, followed by Annexin V/propidium iodide (PI) flow cytometry analysis (**Fig. 4b**). CASCAD-7 treatment for 12 hours produced a pronounced increase in Annexin V-positive apoptotic populations relative to untreated cells and control nanostructures. CASCAD-7 induced significantly higher apoptotic cell fractions compared to CARD-only and CASCAD-0 controls, levels observed with staurosporine treatment (**Fig. 4c**). In contrast, minimal apoptotic activation was detected in untreated cells and CARD-only controls, indicating that efficient apoptosis requires both nanoscale CARD clustering and intracellular scaffold delivery. The apoptotic activity induced by CASCAD-7 was substantially greater, supporting a mechanism in which organized heptameric clustering of CARD domains drives efficient activation of the endogenous caspase cascade through proximity-induced assembly.

We next examined the relationship between CASCAD-7 concentration and apoptotic output (**Fig. 4d**). CASCAD-7 exhibited a concentration-dependent increase in apoptosis up to an intermediate concentration range, after which apoptotic activity plateaued or modestly declined at higher concentrations. The fitting of response curve identified a peak activation at ∼ 5.8 nM CASCAD-7. This behavior mirrors the stoichiometric dependence observed in the biochemical caspase-9 activation assays and requiring optimal number of procaspase-9 molecules onto individual CASCAD-7. Excess scaffold concentrations may distribute procaspase-9 molecules across separate nanostructures, thereby reducing effective intermolecular dimerization and limiting catalytic activation efficiency. Cell viability measurements post 24-hour treatment further confirmed these findings (**Fig. 4e**). CASCAD-7 treatment significantly reduced relative cell viability compared to untreated cells, scaffold-only controls and CARD-only conditions. Importantly, viability loss induced by CASCAD-7 approached levels comparable to staurosporine treatment, indicating that programmable nanoscale organization of CARD domains is sufficient to induce robust apoptotic signaling and downstream cell death.

To extend beyond leukemic cell line and further confirm the mechanistic specificity of CASCAD-7-induced apoptosis, we compared apoptotic responses in prototypical models of liver (HepG2) and breast (MCF-7) cancers using Annexin V/PI analysis. HepG2 and MCF-7 are highly valued for retaining many functions of human hepatocytes and hormone-dependent breast cancer, respectively. CASCAD-7 significantly increased apoptotic populations in HepG2 cells relative to CASCAD-0 controls, whereas apoptotic activation was markedly lower in MCF-7 cells (**Fig. 4f-g** & **Supplementary Fig. 12)**. In contrast, staurosporine treatment produced measurable apoptotic populations in both HepG2 and MCF-7 cells (**Fig. 4f-g**), consistent with its broad pro-apoptotic activity that is not restricted to a single signaling pathway^63^. Because MCF-7 cells are deficient in caspase-3^64^, a critical executioner caspase downstream of caspase-9, the reduced response supports a mechanism in which CASCAD-7 engages the Caspase-9 mediated cascade rather than inducing nonspecific cytotoxicity. The ability of CASCAD-7 to activate apoptosis in caspase-competent cells while exhibiting reduced efficacy in caspase-3-deficient cells hints that scaffold-mediated intracellular signaling is mechanistically linked to endogenous apoptotic machinery.

### CASCAD-7 induces apoptotic signaling in leukemia 3D spheroids

To evaluate whether CASCAD-7 retains apoptotic functionality within a physiologically relevant three-dimensional (3D) microenvironment, we established a 3D leukemia spheroid model using HEL92.1.7 cells encapsulated within PEG 4-arm hydrogels (**Fig. 5a**). Although leukemia cells primarily exist as circulating cells in vivo, forming multicellular aggregates within bone marrow and tissue-associated leukemic niches is associated with metastatic disease and enhanced survival, drug resistance and altered apoptotic signaling^65,66^. 3D leukemia spheroids therefore provide a physiologically relevant model for evaluating both intracellular nanostructure activity within densely packed multicellular leukemic assemblies^67^ and broad applicability of our method to a range of cancer conditions. HEL92.1.7 cells were initially embedded as single cells within the hydrogel matrix and cultured over time to allow spontaneous spheroid formation and growth. Brightfield imaging demonstrated progressive spheroid maturation over a 10-day period, with spheroid diameter increasing substantially between Day 0 and Day 10 (**Supplementary Fig. 13a**). Quantitative analysis confirmed robust spheroid growth kinetics, yielding multicellular aggregates with diameters approaching ∼50-80 µm at later time points (**Supplementary Fig. 13b**). Based on spheroid morphology and size distribution, Day 6-7 spheroids were selected for subsequent experiments because they exhibited relatively uniform aggregate size and structural organization across the culture population.

**Fig. 5:**
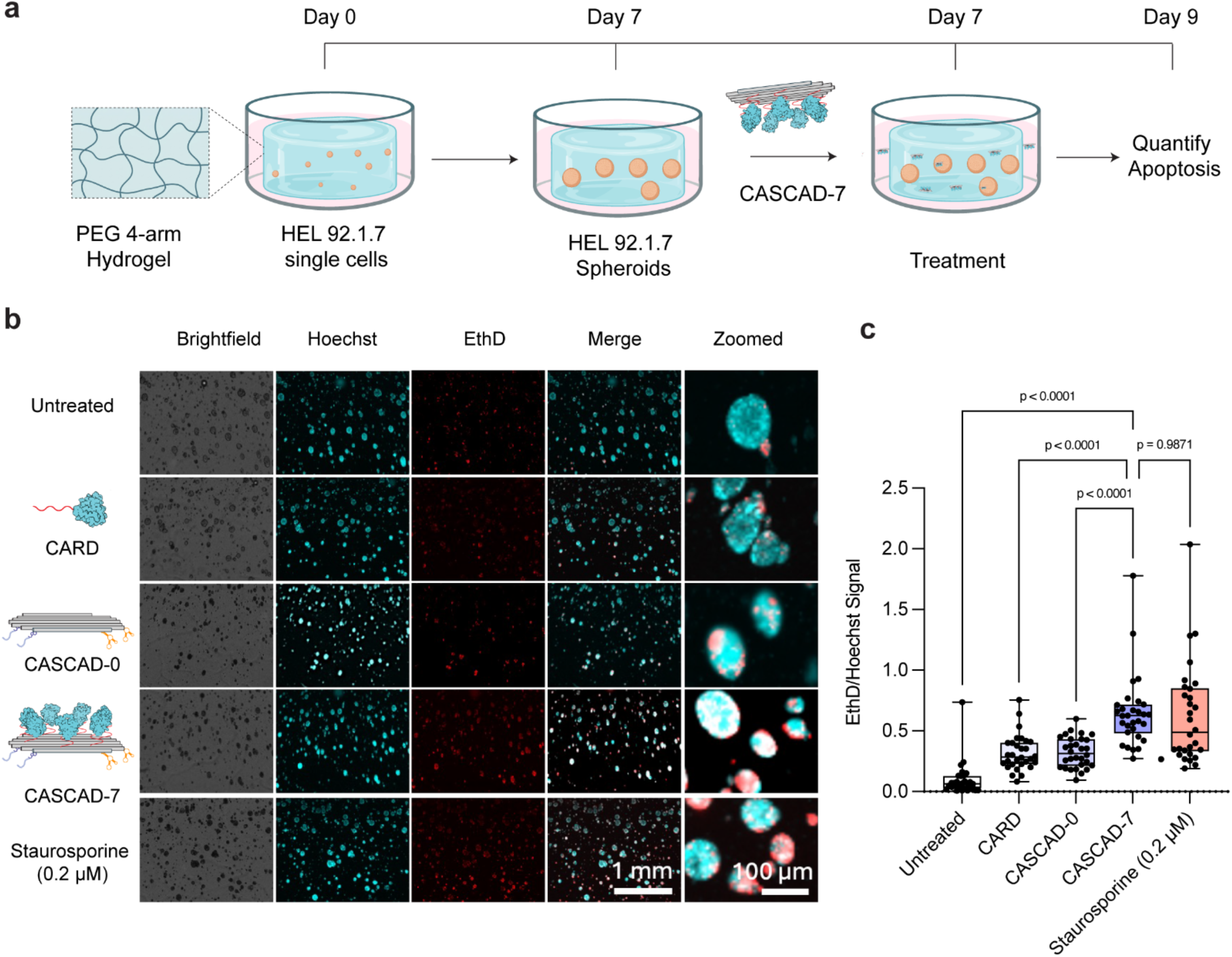
CASCAD-7 induces apoptotic signaling in HEL 92.1.7 leukemia 3D spheroids. **a.** Experimental schematic. HEL 92.1.7 cells were embedded as single cells in PEG 4-arm hydrogel on Day 0 and allowed to form spheroids over 7 days. On Day 7, spheroids were treated with the CASCAD-7 and apoptosis was quantified on Day 9. **b.** Representative fluorescence microscopy images of spheroids under each treatment condition. Nuclei were stained with Hoechst (cyan) and dead/dying cells with Ethidium Homodimer (EthD; red), shown in brightfield, individual fluorescence channels, merged, and zoomed views. **c.** Quantification of apoptosis expressed as the EthD/Hoechst signal ratio for HEL spheroids. CASCAD-7 treatment significantly increased cell death compared to untreated controls, CARD, and CASCAD-0, while no significant difference was observed between CASCAD-7 and Staurosporine (0.2 µM). Each dot represents an individual spheroid. Data are presented as mean ± s.e.m for *n* = 30 spheroids. P values determined by one-way ANOVA followed by Dunnett’s multiple comparisons test.

Following spheroid maturation, cultures were treated with CASCAD-7, CARD-only controls, non-functionalized origami scaffolds (CASCAD-0), or staurosporine (0.2 µM) as a positive apoptosis control (**Fig. 5a**). Apoptotic activity was quantified two days post-treatment using a dual-fluorescence viability assay based on Hoechst nuclear staining and Ethidium Homodimer (EthD), which labels membrane-compromised apoptotic and dead cells. Fluorescence imaging revealed a pronounced increase in EthD-positive regions in CASCAD-7-treated spheroids relative to untreated controls and control nanostructures (**Fig. 5b**). In contrast, untreated spheroids and spheroids treated with either free CARD domains or CASCAD-0 exhibited only minimal EthD incorporation, indicating limited induction of apoptosis under these conditions. Notably, CASCAD-7-treated spheroids displayed EthD staining patterns qualitatively comparable to those observed following staurosporine treatment, suggesting efficient activation of apoptosis within the three-dimensional cellular architecture.

Higher-magnification imaging further demonstrated extensive EthD accumulation throughout CASCAD-7-treated spheroids, indicating that apoptotic activation was not restricted to peripheral cells but extended throughout multicellular aggregates (**Fig. 5b, zoomed images**). This observation is particularly significant given the diffusion barriers and restricted intracellular accessibility typically associated with dense 3D cultures that reconstitute tumor tissues. The ability of CASCAD-7 to induce apoptosis within spheroids therefore suggests that the nanostructure maintains functional intracellular delivery and activity even under spatially constrained microenvironmental conditions. Quantitative image analysis confirmed these observations (**Fig. 5c**). CASCAD-7 treatment resulted in a significant increase in the EthD-to-Hoechst fluorescence ratio relative to untreated cells, CARD-only controls and CASCAD-0-treated spheroids. Importantly, apoptotic activity induced by CASCAD-7 approached levels comparable to those observed with staurosporine treatment, while no statistically significant difference was observed between the two conditions. These findings demonstrate that programmable nanoscale organization of CARD domains within CASCAD-7 is sufficient to robustly activate apoptotic signaling in leukemia spheroids cultured within a biomimetic three-dimensional hydrogel environment.

## CONCLUSION and OUTLOOK

CASCAD represents a unique class of synthetic SMOC that reconstitutes the key structural and functional features of natural apoptosomes through nanoscale organization of CARDs. By engineering a heptameric CARD assembly with defined spacing and geometry, CASCAD-7 enables multivalent recruitment of caspase-9 and produced ∼190-fold increase in binding affinity relative to monomeric CARDs. It further promotes valency-dependent and proximity-induced caspase-9 catalysis, demonstrating that nanoscale organization can directly regulate caspase-9 signaling. Notably, CASCAD-7 features targeting aptamers and membrane-interacting peptides that promotes cellular uptake and cytosolic delivery, facilitating its intracellular functionality. In cancer cells, CASCAD-7 induces robust intracellular caspase-9 activation and programmed cell death, with activity approaching staurosporine-treated controls. Importantly, comparative studies in caspase-3-deficient cells suggest that CASCAD-7 indeed engages the caspase-9 mediated intrinsic apoptotic pathway, validating our desired mechanistic specificity. CASCAD-7 also retained apoptotic functionality in 3D leukemia spheroids cultured within synthetic extracellular matrix hydrogels, demonstrating efficiency in physiologically relevant multicellular environments.

The results of our study also broadly present the design principles for engineering DNA nanomaterial based SMOCs, where signaling activity is governed by the nanoscale organization of modular interaction domains. Despite substantial diversity in biological functions, many SMOCs share common organizational principles in which signaling output is governed by the stoichiometry of interacting moieties. Examples include the apoptosome, inflammasome, myddosome and FADDosome, which utilize homotypic interaction domains such as caspase recruitment domains (CARDs), death domains (DDs) and death effector domains (DEDs) to nucleate higher-order signaling assemblies and drive proximity-induced catalytic activation^68^.

Inspired by these naturally occurring signaling complexes, we envision DNA nanostructures as a programmable framework for engineering synthetic SMOCs with tunable structural and functional properties. Unlike conventional biomaterial systems, DNA nanostructures enable independent control over multiple nanoscale design parameters that can collectively govern supramolecular signaling behavior. These include: (i) the identity of the interacting domain, (ii) domain valency, (iii) intermolecular spacing, (iv) geometric organization, and (v) scaffold stoichiometry (**Fig. 6**). Different SMOCs utilize distinct homotypic interaction modules to recruit downstream effectors, including CARDs in apoptosomes and inflammasomes, DDs in myddosomes and DEDs in death-inducing signaling complexes^69^. By incorporating such modules onto programmable DNA scaffolds, diverse synthetic signaling complexes can in principle be constructed using a common nanoscale engineering strategy. Valency and intermolecular spacing govern multivalent interactions and cooperative activation^2^, as nanoscale separation directly influences productive intermolecular dimerization and catalytic activation. Beyond spacing, geometric organization also plays a major role in supramolecular signaling, with DNA nanotechnology enabling architectures ranging from linear assemblies to disk-like and three-dimensional clustered organizations^70^. Finally, scaffold stoichiometry, including the ratio between interacting modules can be fine-tuned through control of binding-site occupancy to optimize signaling efficiency^71^. Within this framework, CASCAD represents a first implementation of a synthetic DNA nanomaterial-based SMOC capable of programmed intracellular caspase activation. In summary, our study highlights the use of DNA nanotechnology as a modular platform for studying and reconstructing higher-order signaling assemblies, opening opportunities for both mechanistic interrogation of spatial signaling biology and development of next-generation therapeutic systems.

**Fig. 6:**
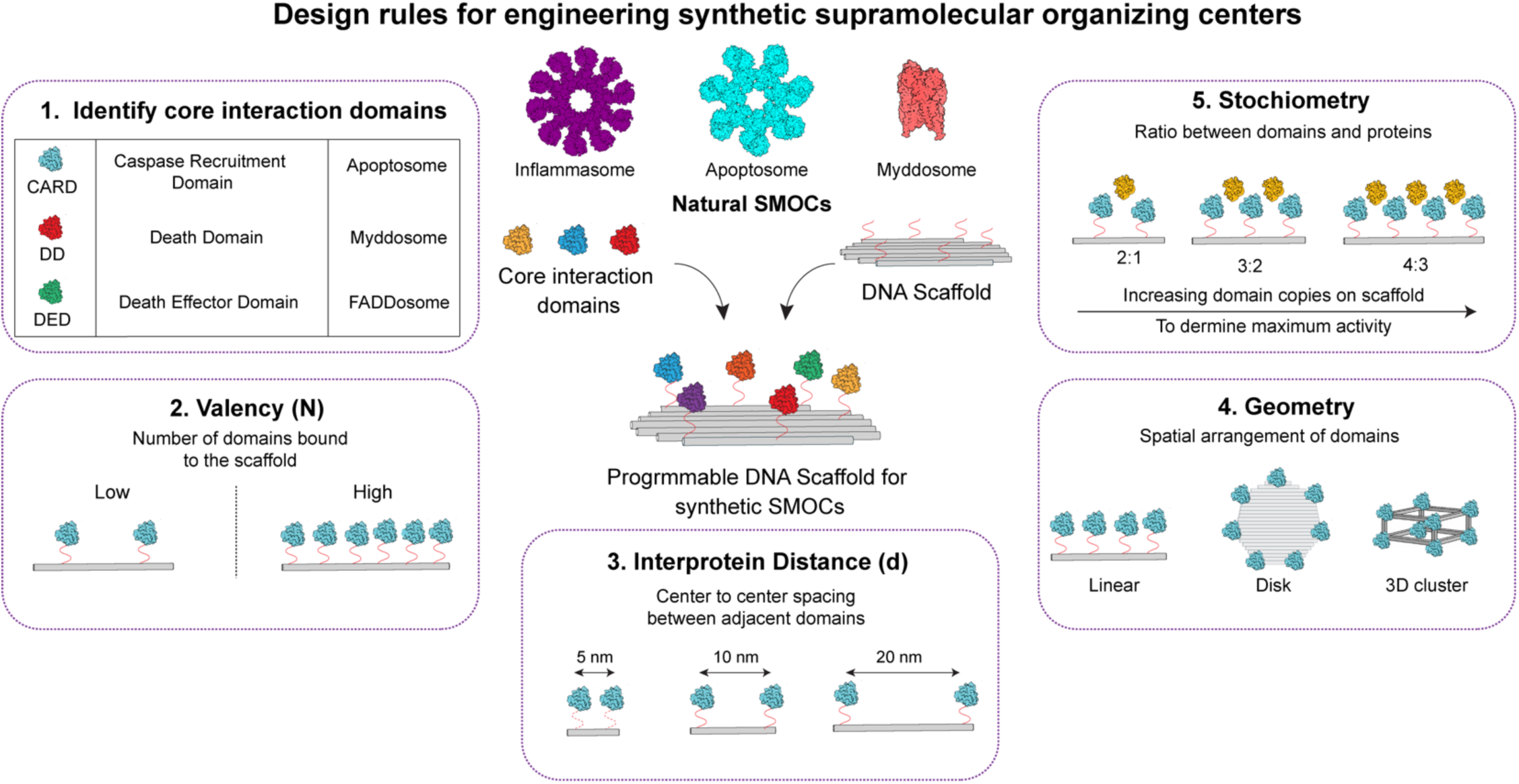
Design rules for engineering synthetic supramolecular organizing centers. Programmable DNA nanostructures provide a modular framework for reconstructing supramolecular organizing centres (SMOCs) through precise control of core interaction domain, valency, intermolecular spacing, geometry and scaffold stoichiometry. Different signaling interaction modules can be spatially organized on DNA origami scaffolds to engineer synthetic intracellular signaling architectures with tunable assembly and activation properties.

## Acknowledgement

We would like to acknowledge the Materials Research Laboratory Central Research Facilities at University of Illinois Urbana-Champaign where AFM images were taken. We also acknowledge the Beckman Institute for Advanced Science and Technology and the Tumor Engineering and Phenotyping Facility of the Cancer Center at Illinois, supported by the National Cancer Institute of the National Institutes of Health (NIH) grant number P30CA275774, where Confocal and fluorescence imaging was performed. This work was supported in part by grant from NSF 2127436 to X.W. R.M. was supported by the National Institute of Biomedical Imaging and Bioengineering of the NIH under Award Number T32EB019944. The content is solely the responsibility of the authors and does not necessarily represent the official views of the National Institutes of Health. R.B. is a Biohub Investigator.

## Notes

### Competing Interest Statement

The authors have declared no competing interest.

